# Effects of a novel *Paraburkholderia* phage IPK on the phenanthrene degradation efficiency of the PAH-degrading strain *Paraburkholderia caledonica* Bk

**DOI:** 10.1101/2025.05.02.651996

**Authors:** Esteban E. Nieto, Nawras Ghanem, Robertina V. Cammarata, Felipe Borim-Correa, Bibiana M. Coppotelli, Antonis Chatzinotas

## Abstract

Phages are a major cause of bacterial mortality, affecting bacterial diversity and ecosystem functioning. However, the impact of phage-host interactions in contaminated environments and their role in pollutant biodegradation have been largely overlooked. We isolated and characterized a novel phage from a polycyclic aromatic hydrocarbon (PAH)-contaminated soil that infects the PAH-degrading bacterium *Paraburkholderia caledonica* Bk, and investigated the effect of different multiplicity of infection (MOI) on the degradation efficiency of phenanthrene. The phage IPK is a temperate phage with a wide pH and temperature tolerance and a burst size of 80 PFC.ml□^1^. The IPK phage was classified as a member of the Caudoviricetes, related to *Pseudomonas* and *Burkholderia* phages; however, its low intergenomic similarity indicates that it belongs to a new species. Three AMGs related to amino acid metabolism and to bacterial growth regulation were identified in the phage genome. The highest multiplicity of infection (MOI 10) showed a rapid recovery of the host density abundance and greater phenanthrene degradation than MOIs ranging from 0.01 to 1. This work highlights the critical role of phage-host interactions in modulating pollutant degradation efficiency, which could be a key for improving the establishment of inoculants in bioremediation processes.

## Introduction

Phages play a key role as a major top-down regulator of bacterial abundance, affecting bacterial diversity and ecosystem functioning. They employ a range of infection strategies, with lytic and lysogenic cycles being the best described lifestyles, although alternative strategies have also been documented (Mäntynen et al. 2021; Chevallereau et al. 2022). Lytic phages infect and replicate in their hosts, releasing the new progeny by killing their hosts. Lysogenic or temperate phages can also enter into a lysogenic cycle in which the phage genome integrates into the host chromosome as a prophage and replicates with the bacterial chromosome. Thus, while lytic phages maintain an antagonistic relationship with their hosts, the fitness of temperate phages and their hosts are more aligned, leading to more mutualistic behaviour (Obeng et al. 2016; Chevallereau et al. 2022; Jurburg et al. 2023). A common outcome of lysogeny is superinfection exclusion, where a prophage protects its host from infection by other closely related phages (Hampton et al. 2020). In addition, temperate phages can broaden their bacterial hosts’ metabolic repertoire by carrying auxiliary metabolic genes (AMGs), promoting the transfer of antibiotic resistance genes, virulence factors, biogeochemical nutrient cycling, and bacterial adaptation and evolution (Zimmerman et al. 2020; Chevallereau et al. 2022).

Several models have been developed to describe phage-host dynamics (Knowles et al. 2016; Silveira et al. 2021; Voigt et al. 2021; Brown et al. 2022). The “Killing-the-winner” (KtW) model proposes that lytic phages suppress fast-growing bacteria, leading to the control of bacterial community abundance and ensuring the coexistence of less competitive species (Thingstad 2000). The “Piggyback-the-winner” (PtW) and “Piggyback-the-loser” (PtL) models describe temperate phage dynamics and focus on the switch from the lytic to the lysogenic cycle as the host density changes, characterised by an increase in lysogeny at high and low bacterial densities, respectively (Knowles et al. 2016; Silveira et al. 2021). In addition, bacterial hosts indirectly control phage production by their physiology, such as growth rate, which is linked to resource availability (Zimmerman et al. 2020).

At the ecosystem level, phages affect soil functioning by being a major cause of bacterial mortality (Kuzyakov and Mason-Jones 2018) and regulate microbial diversity by facilitating the coexistence of species (Voigt et al. 2021; Carreira et al. 2024). Phage activity also affects biogeochemical cycling through the viral shunt, as virus-induced mortality releases nutrients, making them available for uptake by other organisms (Braga et al. 2020; Carreira et al. 2024). Notably, there is some evidence that phages from polluted environments show genomic differences to those from unpolluted environments. Lysogeny and polyvalent strategies tend to be more common in stressful environments (Huang et al. 2024), and an enrichment of AMGs related to stress tolerance and xenobiotic degradation has been reported (Huang et al. 2021; Zheng et al. 2022; Yuan et al. 2023). These genes could benefit the bacterial adaptive response and increase pollutant degradation, but the role of phages in bioremediation still needs to be explored (Ru et al. 2023).

Bioaugmentation, which involves the inoculation of pollutant-degrading microbes, has been proposed as a promising solution for the remediation of contaminated ecosystems (El Fantroussi and Agathos 2005; Muter 2023). However, bacterial inoculants frequently fail to establish, reducing reliability and field success (Kaminsky et al. 2019; Jurburg et al. 2022). An increase in lytic-phage activity may limit the application of microbial-based technologies by reducing inoculum survival, and consequently, the desired metabolic activity (Fu et al. 2009; Albright et al. 2022). Furthermore, the inoculation of allochthonous bacteria, which is a common strategy in biotechnological solutions, may be more affected by the phage community than native bacteria (Braga et al. 2020). However, the role of the phages in controlling inoculum abundance and functionality has received little attention.

In our previous work, we inoculated a polycyclic aromatic hydrocarbons (PAH)-degrading consortium, consisting of the two PAH-degrading strains *Sphingobium* AM and *Paraburkholderia caledonica* Bk, in soils with different pollution exposure histories (Nieto et al. 2024). Inoculation did not increase PAH removal in the chronically contaminated soil, which was explained by the low survival of the inoculated strains due to the response of the eukaryotic predatory community. To further investigate the role of other potential predators in inoculum survival and functioning, this study provides a first characterization of the interaction between a phage and one host strain during PAH-degradation at a laboratory scale. We successfully isolated and characterised a phage from the chronically contaminated soil which infects *P. caledonica* Bk. This strain has the genomic potential for PAH degradation, being able to use these pollutants as sole carbon and energy source (Macchi et al. 2021; Nieto et al. 2023, 2025), and showed lower survival than strain AM in the chronically contaminated soil (Nieto et al. 2024). In addition, we investigated the effect of different phage-bacteria ratios (i.e., the multiplicity of infection (MOI)) on bacterial population dynamics and phenanthrene degradation at the laboratory scale. We hypothesised that a higher initial MOI would result in a lower degradation efficiency due to higher host mortality.

## Materials and methods

### Screening of prophage and CRISPR-Cas arrays in the host genome

The genome of *Parabukholderia caledonica* Bk was previously sequenced (accession number: NHOM01; (Macchi et al. 2021)). We searched for prophage regions in the host genome using the PHASTEST web server (https://phastest.ca/) (Wishart et al. 2023). In addition, CRISPR (clustered regularly interspaced short palindromic repeats) arrays and their associated (Cas) proteins were detected using the CRISPRCasFinder web server (https://crisprcas.i2bc.paris-saclay.fr/CrisprCasFinder/Index) (Couvin et al. 2018).

### Phage enrichment, isolation, purification

A chronically contaminated soil (IPK) from a petrochemical plant in Ensenada, Argentina (34°53’19”S, 57° 55’ 38”W) was selected to isolate phages for the allochthonous PAH-degrading strain *P. caledonica* Bk. The IPK soil was previously treated by landfarming, with several applications of petrochemical sludge. Sampling was done approximately 10 years after the cessation of the petrochemical sludge treatments, showing a total PAH concentration of 573± 138 mg. kg---^-1^ dry soil (Cecotti et al. 2018; Festa et al. 2024).

For phage enrichment, 10 g of soil was mixed with 90 ml of LB broth supplemented with CaCl_2_ and CaSO-_4_ (1 mM) and 1 ml of an overnight culture of the Bk strain. After 24 h of incubation, 5 ml of each enrichment culture was centrifuged for 10 min at 3600 rpm, and the supernatant was filtered through a 0.22 μm nylon membrane. The filtrate was tested for phage activity against *P. caledonica* Bk using a double agar assay with 0.6% soft top agar. Plaques were collected with a sterile pipette tip and transferred to 200 μl of LB broth, incubated for 1 h at room temperature, and centrifuged for 10 min at 15000 g at 4°C. The supernatant was filtered (0.22μm) and tested for phage activity. This process was repeated twice to ensure pure isolates.

### One-step growth curve

The one-step growth curve of the phage was determined by using a modified protocol previously described (Chen et al. 2020). Briefly, 5 ml of an overnight bacterial culture (OD_600_ of 0.3 to 0.5) was centrifuged at 8,000 rpm for 5 min. The cell pellets were resuspended in 500 μl of LB medium, and infected with 100 μl of phage suspension to yield a multiplicity of infection (MOI) of 0.01. After adsorption for 10 min at room temperature, the phage-host mixtures were centrifuged at 12,000 rpm for 10 min to remove unadsorbed phage particles. The cell pellets were resuspended in 5 ml of LB medium and incubated at 30°C with constant shaking. Aliquots were collected every 20 min for up to 3 h and were immediately serially diluted. Phage titers were determined using the spotting plaque assay technique. Three independent replicates were performed for each assay.

### Killing curve

Cultures of *P. caledonica* Bk were infected at an early exponential phase (OD_600_ of 0.4) with different phage concentrations to obtain final MOIs of 0.001, 0.01, 0.1, and 1. Positive and negative controls were also prepared by excluding the phage or bacterial cells, respectively. After incubation in a microplate reader for 13 h at 30 °C, the OD_600_ of each well was measured at 30 min intervals after mixing of 10 sec, with four replicates for each treatment.

### Thermal and pH stability assays

For the thermal stability test, 500 μl of filter-sterilised phage samples were incubated at 15°, 30°, 40° 50°, 60° and 70°C for 24h. For the pH stability test, 40 μl of phage lysate were added to 3960 μl TM buffer with a pH ranging from 1 to 13 and incubated for 1h and 24 h. After incubation, the phage titers were determined by a double-layer assay to determine the number of plaque-forming units.

### Phage DNA extraction, genome sequencing and assembly

The phage DNA was extracted from the lysates, as described by (Thurber et al. 2009). The isolated phage DNA was sequenced using the Oxford Nanopore MinION platform (MN45708). Library preparation followed the “Ligation Sequencing gDNA - Native Barcoding Kit 24 V14 Oxford” protocol (v. NBE_9169_v114_revH_15Sep2022) from Oxford Nanopore Technologies, with one modification: during the DNA repair step, incubation was extended to 15 minutes at 20°C and 65°C. Approximately 10 ng/μl of DNA was pooled to create a barcoded library with a final volume of ∼50 μl, which onto an R10.4.1 (FLO-MIN114) flow cell in accordance with manufacturer instructions. Sequencing runs were conducted and monitored through the MinKNOW software. The raw signal data were first basecalled using Guppy basecaller (v. 6.5.7+ca6d6af). The input fast5 files were processed using the high accuracy (HAC) basecalling model (dna_r10.4.1_e8.2_400bps_hac.cfg), optimized for the R10.4.1 flow cell chemistry.

Demultiplexing was conducted during the basecalling process using the barcoding scheme provided in the Native Barcoding Kit 24 (SQK-NBD114-24). To ensure all reads were retained for analysis, quality score filtering was disabled. To improve the accuracy of basecalling automatic calibration detection was enabled. Guppy was also employed for demultiplexing. Adapter sequences and other residual non-biological elements were trimmed using Porechop (v.0.2.3_seqan2.1.1) (Wick et al. 2017).

Genome assembly was performed using Flye (v.2.7-b1585) (Kolmogorov et al. 2019), which produced high-quality draft genomes with sufficient contiguity for subsequent error correction steps. The draft assemblies were further refined using Medaka (v.1.8.0) (Oxford Nanopore Technologies 2018). For prophage validation, CheckV (v.1.0.3) (Nayfach et al. 2021) and VirSorter (v.2.2.4) (Guo et al. 2021) were utilized. CheckV also assessed the quality and completeness of the prophage sequences. To assign taxonomic classifications, the assembled genomes were analyzed using GTDB-Tk (version v.2.3.2), a software tool based on the Genome Taxonomy Database (GTDB). Automatic annotation was performed using geNomad (v.1.7.1) and VIBRANT (v.1.2.1). Additionally, the unclassified proteins were manually classified using HHpred (probability > 0.9, e-value < 10^−5^) (Söding et al. 2005)

### Phylogenetic and comparative genomic analyses

To elucidate the taxonomy of *Paraburkholderia* phage IPK, most related phages were identified using the NCBI Virus database (https://www.ncbi.nlm.nih.gov/labs/virus/vssi/#/), vConTACT2 (Jang et al. 2019) and VipTree (https://www.genome.jp/viptree/) (Nishimura et al. 2017). Nine complete genomes were obtained from the NCBI dataset, and only two genomes were cluttered according to vContact2 (accession numbers: EU982300.1; OK665841.1). From the VipTree results, we selected 20 closely related phage genomes using the S_G_ score (Zhu et al. 2024). From these genomes, we constructed a phylogenetic tree using the Genome-BLAST Distance Phylogeny (GBDP) method in the Virus Classification and Tree Building Online Resource (VICTOR) (Meier-Kolthoff and Göker 2017). The resulting intergenomic distances were used to infer a balanced minimum evolution tree with branch support via FASTME including SPR postprocessing for the formulas D0. Additionally, intergenomic similarities were calculated using VIRIDIC (Moraru et al. 2020).

### Effect of multiplicity of infection (MOI) on phenanthrene (PHN) degradation

To evaluate the effect of different phage-host ratios (i.e. multiplicity of infection (MOI)) during PHN degradation, *P. caledonica* Bk was grown in LB medium overnight at 30 °C and 150 rpm, centrifuged at 6000 rpm for 10 min, then washed three times with 0.85% NaCl and resuspended in the same solution. A density of 1*10^6^ CFU.ml^-1^ was inoculated into 10 ml of Liquid Mineral Medium (LMM) (Vecchioli et al. 1990) supplemented with 200 mg.l^-1^ of PHN as the sole carbon and energy source. Since no reduction in host OD was observed in the killing curve at the two lowest MOIs, we excluded the MOI 0.001, added a higher one and tested the following MOIs: 0.01, 0.1, 1, and 10. Bk cultures without phages were used as control. Each treatment was carried out by destructive triplicate monitoring at 0, 1, 2, 3 and 4 days of incubation at 30°C and 150 rpm. Three consecutive chemical extractions with ethyl acetate were performed and PHN were measured by HPLC (Waters® XBridge C18 3.5 μm, 4.6 × 65 mm) following (Nieto et al. 2023).

In parallel, triplicate cultures with the different MOIs were run and resampled at 0, 1, 2, 3 and 4 days of incubation to determine phage titers using the spotting plaque assay technique and bacterial abundances after plating on LB agar, and to describe virus-host-ratios over time. Day 4 phage samples were lost during processing so they are excluded in the article.

### Statistical analysis

Statistical analyses were performed in R v 4.3.3(R Core Team, 2024). Two-ways ANOVA were carried out to test the effects of MOI in degradation efficiency with MOI and time as independent variables, using the *rstatix* (v.0.7.2) (Kassambara 2023) R package. Shapiro and Levene tests were performed to check normality and homoscedasticity, respectively. Tukey’s Honestly Significant Difference (HSD) post-hoc test was used to perform pairwise comparisons between group means. Due to the lack of normality, significant differences between log_10_ of bacterial abundance across different MOIs over time were assessed using non-parametric Kruskal-Wallis test and Dunn test as a post-hoc test to perform pairwise comparisons between group means. PHN concentration, bacterial and phage counts are expressed as mean ± standard deviation.

## Results

### Screening of prophage and CRISPR-Cas arrays in the host genome

No prophage sequences were found in *P. caledonica* Bk. Three CRISPR elements were identified that were not associated with *cas* genes; the low level of evidence (level 1) indicates that the arrays are invalid (Couvin et al. 2018).

### Biological characterization of phage

One phage was isolated from the chronically PAH-contaminated soil for the host *P. caledonica* Bk and was named *Paraburkholderia* phage IPK. Thermal and pH stability tests were performed and showed survival in a pH range of 4-11 and thermal stability up to 60 °C (Figure 1). The one-step growth curve showed that phage IPK had a latent period of 80 min and a burst size of 80 PFU.cell^-1^ (Figure 1C). The effect of MOI on *P. caledonica* Bk survival was assessed using four MOIs (0.001, 0.01, 0.1 and 1). The lower MOIs did not show inhibition of the host growth during the analysed period, whereas both MOIs of 0.1 and 1 inhibited growth. The highest MOI showed the largest inhibition, reducing growth within five hours of incubation (Figure 1D).

**Figure 1:**
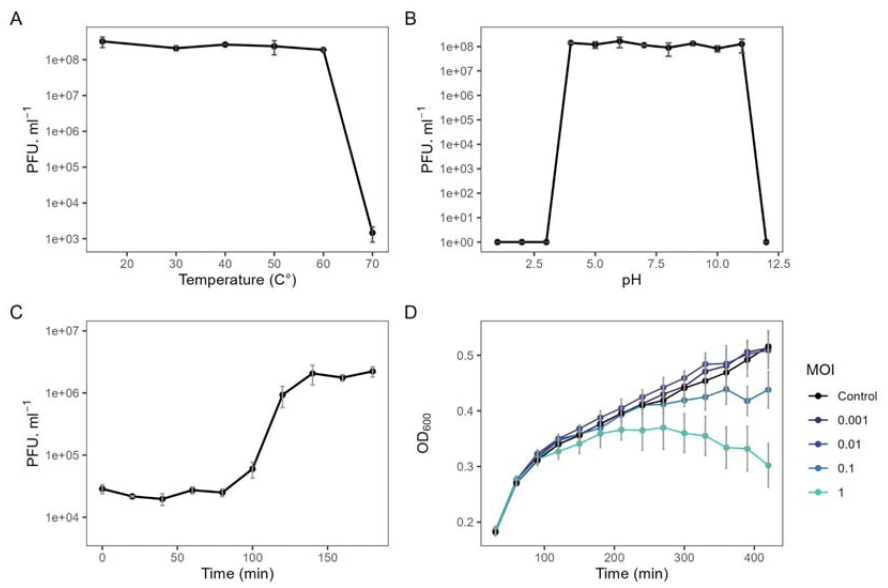
Biological characterization of phage IPK. A) Thermostability profile. B) pH stability profile. C) One-step growth curve using an MOI of 0.01. D) Killing curve at different MOI.

### Phage genome characterization

Whole genome analysis showed that phage IPK is a dsDNA phage with a genome size of 40,356 bp and a GC content of 60.42%. A total of 63 ORFs were predicted, of which 43 were functionally annotated while 20 were annotated as hypothetical phage proteins (Figure 2, Table S1). Around 25% were classified as structural proteins, including spike proteins, tail tube and tail sheath proteins, and putative major capsid proteins. Lifestyle prediction showed that the phage was temperate (99.98% probability). Among the ORFs related to life cycle, an integrase and an excisionase were annotated, which facilitate phage genome integration or excision into the host genome, respectively. In addition, an antirepressor protein KilAC and a phage regulatory protein CII were annotated in the genome, which play a role in the regulation of the lysogenic cycle. Three ORFs were annotated as auxiliary metabolic genes, including the antitoxin HicB and the mRNA interferase toxin HicA, which are involved in the regulation of the regulation of the bacterial growth, and a cysteine dioxygenase, which is involved in the metabolism of cysteine.

**Figure 2:**
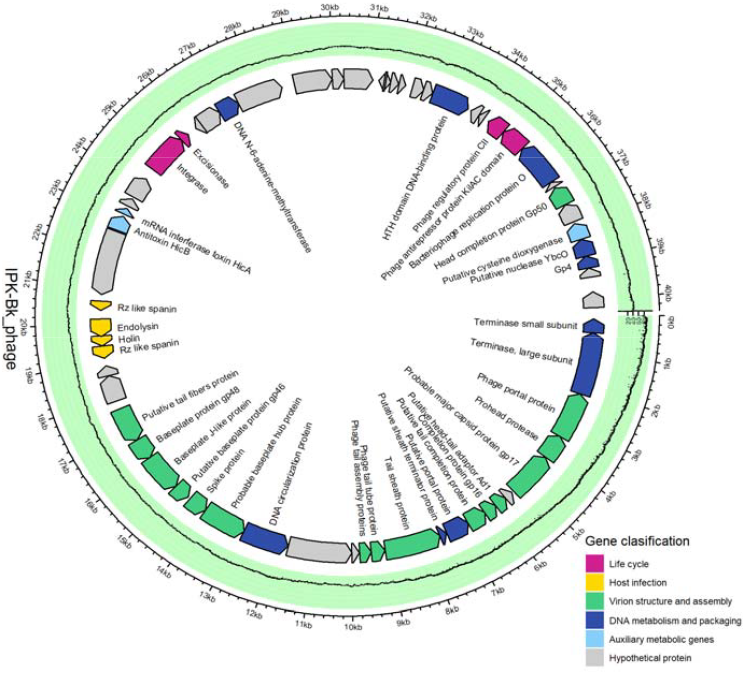
Genome overview of phage IPK. The different colours indicate different types of functional modules

### Phylogenetic and comparative genome analysis

The genomic analysis revealed that the phage IPK belongs to the class *Caudoviricetes*. To obtain a more detailed classification, a phylogenetic analysis was performed using the VICTOR method, focusing on the phages with the highest similarity identified by the NCBI Virus database, vContact2 and VipTree (Figure 3A). According to these results, phage IPK clustered with other *Burkholderia* and *Pseudomonas* phages and was more closely related to *Burkholderia* phage BgVeeders33 and *Pseudomonas* phage DVM 2008. The intergenomic similarity analysis showed low similarity with the selected genomes (Figure 3B). Therefore, both analyses suggest that the phage IPK belongs to a new genus.

**Figure 3:**
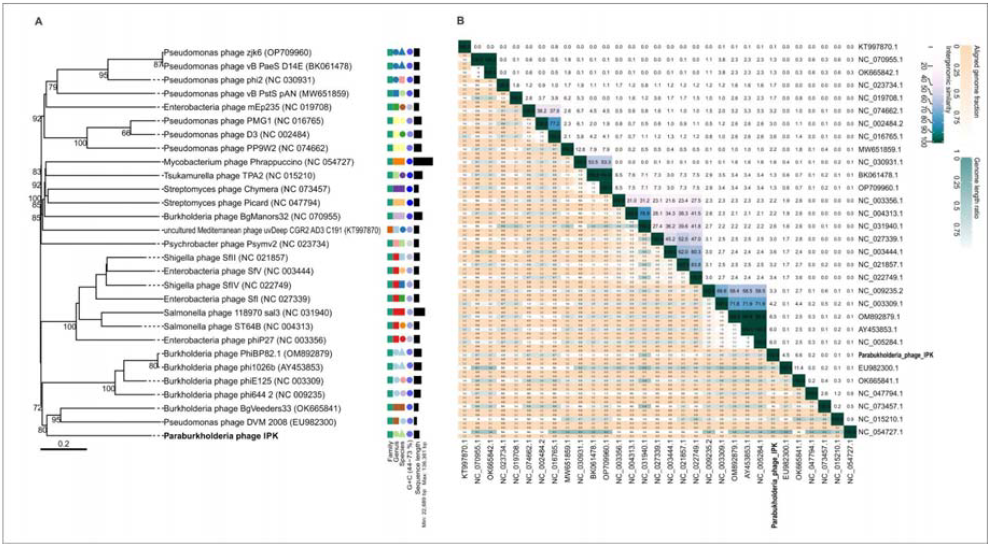
A) Phylogenomic Genome-BLAST Distance Phylogeny (GBDP) method tree inferred using formula D0. The branch lengths of the resulting VICTOR trees are scaled according to the respective distance formula used. B) Heatmap generated by VIRIDIC showing the intergenomic similarity values (right half) and alignment indicators (left half). The percent identity between two genomes was determined by BLASTn, integrating intergenomic similarity values with data on genome lengths and aligned genome fractions.

### Effects of phage density on phenanthrene biodegradation

The MOI affected the biodegradation efficiency of *P. caledonica* Bk (Figure 4). A latency period was observed at all MOIs; however, its duration varied depending on the treatment. After two days of incubation, the control and MOI 10 showed 38.5 ± 4.2% and 15.8 ± 6.2% of PHN degradation, respectively, while no degradation was observed at MOI 0.01, MOI 0.1 and MOI 1. On day three, the latter treatments showed 13.4±11.9%, 32.7 ±10.7% and 18.6 ± 5.4% of degradation, respectively, which were lower (p-value<0.05) than the degradation shown at MOI 10 (51.8± 7.6%) and MOI 0 (76.8± 10.9%). At the end of the incubation period, the highest and lowest degradation were obtained in the control (93.0±7.1%, p-value<0.05) and at MOI 0.1 (30.8± 20.6%), respectively, while the other treatments showed intermediate degradation efficiencies.

**Figure 4:**
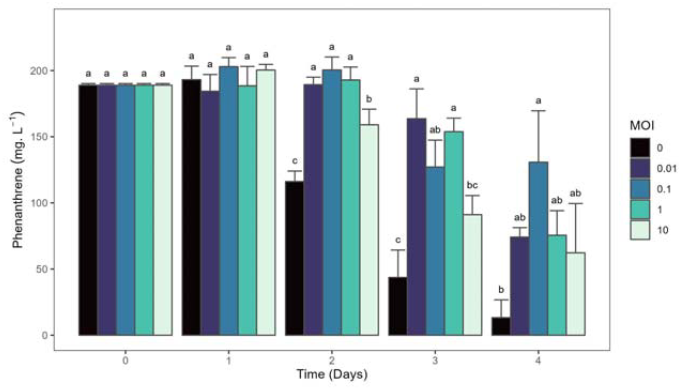
Concentration of phenanthrene (PHN) over time in *P. caledonica* Bk cultures at different MOI (0, 0.01, 0.1, 1 and 10) grown in PHN as only carbon and energy source (200 mg.ml^-1^). The results are expressed as the mean values of independent triplicates and the error bars indicate standard deviation.

The population dynamics of both host and phage were analysed during PHN degradation (Figure 5 A and B). The control showed an increase in the bacterial density from the beginning of the incubation period until day 2, reaching a density of 2.2*10^8^ ± 4.7*10^7^ CFU.ml^-1^, which remained in the same order until the end of the incubation period (Figure 5A). At MOI 0.01, the density of the host remained constant until the end of the incubation period, when an increase was observed. At MOI 0.1, MOI 1 and MOI 10 treatments, the bacterial density decreased compared to the control (*p* < 0.05) and reached values close to 10^5^ CFU.ml^-1^. However, the dynamics in the systems changed after day 3, when MOI 10 showed no difference from the control (∼10^8^ CFU.ml^-1^, *p* >0.05), while similar bacterial abundances were reached at the intermediate MOI treatments at the end of the incubation period (Figure 5A). All treatments showed an increase in phage particles after one day of incubation, with MOI 0.1, MOI 1 and MOI 10 resulting in phage densities close to 10^8^ PFU.ml^-1^(Figure 5B) The MOI 0.01 showed lower phage densities but the highest production rate, with an increase of 4 orders of magnitude compared to the initial density. After three days all treatments showed similar PFU values (Figure 5B).

**Figure 5:**
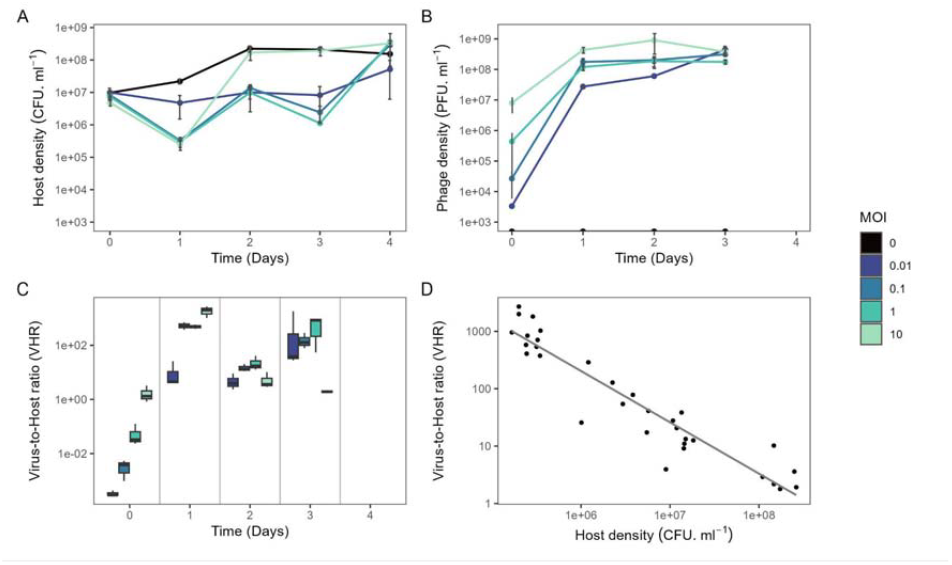
A) bacterial, B) phage counts and C) Virus-host-ratios (VHR) over time in *P*. *caledonica* Bk cultures at different MOI (0, 0.01, 0.1, 1 and 10) grown in PHN as only carbon and energy source (200 mg.ml^-1^). The results are expressed as the mean values of independent triplicates and the error bars indicate standard deviation. D) Virus-to-host ratio as a function of host density

Different virus-host-ratios over the time were observed across the treatments (Figure 5C). Compared to the initial proportion, after three days of incubation the MOI 10 treatment showed a ratio close to 1, whereas in the other treatments, the proportion of phages exceeded that of the bacterial host by two orders of magnitude. The differences between the treatments correlated with the increase in host density observed at MOI 10 (Figure 5A), as the number of PFU.ml^-1^ remained constant (Figure 5B). In addition, a negative correlation was observed between the log of the virus-host-ratio and the density of the host (*R*^*2*^= 0.89, p-value < 0.01, Figure 5D).

## Discussion

Bioaugmentation is considered as a promising and sustainable solution for the remediation of contaminated soils. However, the outcome of inoculation is variable, mainly due to the lack of establishment of the inoculum, which reduces applicability (Kaminsky et al. 2019; Jurburg et al. 2022). After inoculation, the abundance of the inoculated bacteria can trigger potential predators, including phages, to become active, limiting inoculum survival and its biotechnological applicability (Shapiro and Kushmaro 2011; Albright et al. 2022). Although phages can be one of the major causes of bacterial mortality, the ecological implications of phage-mediated top-down regulation of inoculants have received little attention. To our knowledge, only one previous report showed a failure on bioaugmentation due to the lack of establishment of the inoculant as a consequence of phage activity (Fu et al. 2009).

In our previous work (Nieto et al. 2024), the inoculation of a PAH-degrading consortium consisting of the strains *Sphingobium* AM and *Paraburkholderia caledonica* Bk failed to remove PAHs in a chronically PAH-contaminated soil. We tracked the fate of the C^13^-labelled biomass of the consortium using DNA-SIP and identified a rapid response of the eukaryotic predator community after inoculation, which correlated with the low survival of the inoculated strains. However, this work did ot assess the role of phages, which may also contribute to the observed low-inoculum survival. Here, we provide a first description of the host-phage interaction during pollutant degradation by isolating a new phage from the same contaminated soil that infects one of the inoculated strains (*Paraburkholderia caledonica* Bk) and by characterising the effects of different phage-host ratios (i.e. MOIs) on this process under controlled laboratory conditions.

The genus *Paraburkholderia* includes mainly environmental species with a promising biotechnological potential (Vio et al. 2020), whose members show a high frequency of prophages in their genomes, correlating with the diversification of the genus (Pratama et al. 2018). The Bk strain did not have any valid CRISPR-Cas arrays or prophage sequences in its genome, lacking these mechanisms of host immunity against phage infection (Hampton et al. 2020). The isolated *Parabukholderia* phage IPK exhibited a wide range of temperature and pH stability (Figure 1A and B). Phylogenetic analysis suggested that the phage IPK belongs to a new genus related to *Burkholderia* and *Pseudomonas* prophages (DeShazer 2004; Mavrodi et al. 2009; Khrongsee et al. 2024) (Figure 3). The presence of lysogeny-related genes indicates that the isolated phage is a temperate phage. Lysogeny may represent an adaptive strategy for phages to cope with adverse environments (Huang et al. 2024). Some recent studies showed that the number of prophages correlated with contamination severity (Huang et al. 2021; Zheng et al. 2022; Yuan et al. 2023). However, Xia et al (2023) reported that there was a higher proportion of lysogenic phages at low benzo[a]pyrene exposure than at high exposure. Despite variable phage-bacterium interactions, phage-encoded AMGs associated with microbial antioxidant and pollutant degradation were enriched in these contaminated environments, suggesting that phages may play a role in the adaptive response of the host by increasing the biodegradation potential (Zheng et al. 2022; Xia et al. 2023). The genome of phage IPK contained three AMGs, including a cysteine dioxygenase, the antitoxin HicB and the mRNA interferase toxin HicA (Figure 2, Table S1). Xia et al (2023) reported that the AMGs potentially involved in acid metabolism were the most abundant AMGs in lysogenic phages under benzo[a]pyrene exposure, and showed the expression of these AMGs at low exposure levels. In addition, the toxin-antitoxin systems HicAB contribute to the regulation of bacterial growth under stress conditions and can also contribute to the maintenance of the prophage in the host (Qian et al. 2022; Encina-Robles et al. 2024). Therefore, these AMGs may expand the host metabolic profile and enhance microbial environmental adaptability (Huang et al. 2021).

The relationship between phage activity and hydrocarbon degradation is not well understood (Ru et al. 2023). Our results showed that the phage-host dynamics affected the PHN degradation efficiency of the Bk strain (Figure 4). All treatments showed a lag period in PHN degradation, including the control without phages as previously reported (Nieto et al. 2023), but the duration of this period differed between the treatments and correlated with host bacterial density (Figure 5A). After one day, the three highest MOI showed a similar reduction in host density; however, MOI 10 rapidly recovered to densities similar to the control after two days of incubation, when both treatments showed significant PHN degradation. This recovery could be linked to the nutrient released after lysis supporting additional growth and improving biodegradation efficiency (i.e., the viral shunt; Suttle 2007; Rosenberg et al. 2010). In contrast, the other treatments recovered slowly, with no PHN degradation until day three. The degradation latency in Bk has been linked to a delay in the expression of PAH-degrading catabolic genes (Nieto et al. 2023). Other studies have suggested a density-dependent regulation of these genes via quorum sensing in different microbes (Yong and Zhong 2013; Yu et al. 2020). Thereby, a lower degradation efficiency may be a consequence of a lower host density.

Based on these results we reject our hypothesis, as higher initial MOI did not cause a lower degradation efficiency. Higher phage abundance can lead to a higher selective pressure for resistant forms and may contribute to an increase in host abundance (Koskella and Brockhurst 2014), but similar mutation rates should be expected across the treatments, and would not explain the fast recovery observed only at MOI 10. We speculate that an increase in lysogeny, the frequency of which correlates with coinfection rates and higher MOI (Herskowitz and Hagen 1980), and which may confer immunity to the host (Hampton et al. 2020), may be responsible for the rapid recovery and PHN degradation observed at MOI 10. The presence of lysogenic ORFs in phage IPK, in addition to its phylogenetic relationship with other prophages, supports this assumption. Furthermore, we observed a negative correlation between virus-to-host ratio and bacterial abundance (Figure 5D), which is consistent with the pattern previously described both at genus (Coutinho et al. 2017) and community levels (Knowles et al. 2016; Silveira et al. 2021). Based on lysogeny markers, these studies showed a higher frequency of lysogeny at higher host abundances as described by the Piggyback-the-Winner model (Knowles et al. 2016). Further experiments testing the expression of lysogenic markers or the lysogenization of surviving hosts are needed to confirm our interpretation.

Currently, advances in viral metagenomics have improved our understanding of phage diversity. However, the role of phages in host functionality, particularly in polluted environments, remains largely overlooked. Laboratory models continue to be valuable tools for gaining new insights into the complexity of these interactions (Puxty and Millard 2023), which could be crucial for optimising inoculum screening and increasing the applicability of bioaugmentation. The initial phage-host ratio strongly influences both bacterial survival and degradation efficiency. High initial phage pressure can lead to significant bacterial mortality, however, over time, resource release caused by lysis supports bacterial regrowth and accelerates degradation. Interestingly, under such high phage pressure, the potential for lysogeny or phage mutation may offer a selective advantage by stabilizing bacterial populations, allowing host survival in the face of phage predation (Paul, 2008, Weitz et al., 2012). Our findings provide important insights into optimizing phage-host interactions for enhanced bioremediation efficiency, with implications for environmental cleanup strategies.

## Conclusion

Our work demonstrated the effect of a phage-host ratio on the degradation performance of *P. caledonica* Bk, which correlated with host abundance. Notably, the higher initial MOIs showed faster degradation compared to the lower initial MOIs. Although this interaction may change in a complex, structured system such as soil (Koskella et al. 2022), these results highlight the need to consider the role of phage-host interaction on inoculum survival and therefore its function (e.g. degradation efficiency), which may be key to improving inoculation success.

## Data availability

The Paraburkholderia phage IPK genome sequence was deposited in NCBI under the accession number PV588664.

## Author contributions

EEN contributed to the study conception and design, material preparation. Data collection, and analysis were performed by EEN, NG, RVC and FB-C. Supervision was performed by AC and BMC. The first draft of the manuscript was written by EEN and all authors contributed to manuscript editing. All authors read and approved the final manuscript.

## Acknowledgements

EEN received funding by the Research Grants — Short-Term Grants programme of the Deutscher Akademischer Austauschdienst (DAAD) and doctoral fellowship by CONICET. FB-C received funding by the Deutsche Forschungsgemeinschaft (DFG, German Research Foundation)—SFB 1076—Project Number 218627073 as part of the Collaborative Research Centre AquaDiva of the Friedrich Schiller University Jena.

## Ethical approval

Not applicable.

## Consent to participate

Not applicable.

## Consent to publish

Not applicable.

