## Supplemental material for "Effects of a novel *Paraburkholderia* phage IPK on the phenanthrene degradation efficiency of the PAH-degrading strain *Paraburkholderia caledonica* Bk"

1 Centro de Investigación y Desarrollo en Fermentaciones Industriales, CINDEFI (UNLP; CCT-La Plata, CONICET), Street 50 N°227, 1900 La Plata, Argentina

2 Department of Applied Microbial Ecology, Helmholtz Centre for Environmental Research—UFZ, Leipzig, Germany,

3 Institute of Biology, Leipzig University, Leipzig, Germany

4 German Centre for Integrative Biodiversity Research (iDiv) Halle-Jena-Leipzig, Germany.

Table S1: Predicted ORFs, their positions on the genome, size, annotations and probable role in phage life cycle.

| **ORF** | **Start** | **End** | **Length** | **Strand** | **Protein annotation** | **Role/function in phage life cycle** |
| --- | --- | --- | --- | --- | --- | --- |
| ORF1 | 84 | 464 | 381 | + | Terminase small subunit | DNA metabolism and packaging |
| ORF2 | 461 | 2161 | 1701 | + | Terminase large subunit | DNA metabolism and packaging |
| ORF3 | 2161 | 3435 | 1275 | + | Phage portal protein | Virion structure and assembly |
| ORF4 | 3425 | 4108 | 684 | + | Prohead protease | Virion structure and assembly |
| ORF5 | 4120 | 5361 | 1242 | + | Probable major capsid protein gp17 | Virion structure and assembly |
| ORF6 | 5416 | 5637 | 222 | + | Hypothetical protein |  |
| ORF7 | 5634 | 5966 | 333 | + | Putative head-tail adaptor Ad1 | Virion structure and assembly |
| ORF8 | 5968 | 6309 | 342 | + | Completion protein gp16 | Virion structure and assembly |
| ORF9 | 6306 | 6827 | 522 | + | Putative tail completion protein |  |
| ORF10 | 6824 | 7444 | 621 | + | Putative portal protein |  |
| ORF11 | 7454 | 7639 | 186 | + | Putative sheath terminator protein | DNA metabolism & packaging |
| ORF12 | 7636 | 9120 | 1485 | + | Mu-like prophage FluMu tail sheath protein | Host infection |
| ORF13 | 9131 | 9490 | 360 | + | Phage tail tube protein | Virion structure and assembly |
| ORF14 | 9487 | 9792 | 306 | + | Phage tail assembly chaperone proteins | Virion assembly and assembly |
| ORF15 | 9804 | 9998 | 195 | + | Hypothetical protein |  |
| ORF16 | 9991 | 11673 | 1683 | + | Hypothetical protein |  |
| ORF17 | 11676 | 12896 | 1221 | + | DNA circularization protein | DNA metabolism and packaging |
| ORF18 | 12893 | 14050 | 1158 | + | Probable baseplate hub protein | Virion structure and assembly |
| ORF19 | 14050 | 14565 | 516 | + | Spike protein | Host infection |
| ORF20 | 14625 | 15065 | 441 | + | Putative baseplate protein gp46 | Virion structure and assembly |
| ORF21 | 15058 | 16104 | 1047 | + | Baseplate J-like protein | Virion structure and assembly |
| ORF22 | 16095 | 16688 | 594 | + | Putative baseplate protein gp48 | Virion structure and assembly |
| ORF23 | 16688 | 17644 | 957 | + | Putative tail fibers protein | Host infection |
| ORF24 | 17872 | 18591 | 720 | - | Hypothetical protein |  |
| ORF25 | 18594 | 18872 | 279 | - | Hypothetical protein |  |
| ORF26 | 19075 | 19419 | 345 | + | Rz-like spanin | Host infection |
| ORF27 | 19416 | 19688 | 273 | + | Holin | Host infection |
| ORF28 | 19669 | 20148 | 480 | + | Endolysin | Host infection |
| ORF29 | 20380 | 20637 | 258 | + | Rz-like spanin | Host infection |
| ORF30 | 20812 | 22578 | 1767 | + | Hypothetical protein |  |
| ORF31 | 22610 | 23026 | 417 | - | Antitoxin HicB | Auxiliary metabolic genes |
| ORF32 | 23084 | 23260 | 177 | - | mRNA interferase toxin HicA | Auxiliary metabolic genes |
| ORF33 | 23326 | 23553 | 228 | - | Hypothetical protein |  |
| ORF34 | 23613 | 24209 | 597 | - | Hypothetical protein |  |
| ORF35 | 24518 | 25612 | 1095 | - | Integrase | Life cycle |
| ORF36 | 25609 | 25845 | 237 | - | Excisionase | Life cycle |
| ORF37 | 26090 | 26296 | 207 | + | Hypothetical protein |  |
| ORF38 | 26279 | 26857 | 579 | - | Hypothetical protein |  |
| ORF39 | 26854 | 27408 | 555 | - | DNA N-6-adenine-methyltransferase (Dam) |  |
| ORF40 | 27405 | 28634 | 1230 | - | Hypothetical protein |  |
| ORF41 | 28962 | 29960 | 999 | - | Hypothetical protein |  |
| ORF42 | 29962 | 30270 | 309 | - | Hypothetical protein |  |
| ORF43 | 30267 | 31025 | 759 | - | Hypothetical protein |  |
| ORF44 | 31184 | 31321 | 138 | + | Hypothetical protein |  |
| ORF45 | 31364 | 31492 | 129 | - | Hypothetical protein |  |
| ORF46 | 31492 | 31701 | 210 | - | Hypothetical protein |  |
| ORF47 | 31716 | 31892 | 177 | - | Hypothetical protein |  |
| ORF48 | 32074 | 32382 | 309 | - | Hypothetical protein |  |
| ORF49 | 32375 | 32629 | 255 | - | Hypothetical protein |  |
| ORF50 | 32633 | 33655 | 1023 | - | Putative HTH domain DNA-binding protein |  |
| ORF51 | 33738 | 33977 | 240 | + | Hypothetical protein |  |
| ORF52 | 33977 | 34141 | 165 | + | Hypothetical protein |  |
| ORF53 | 34279 | 34758 | 480 | + | Phage regulatory protein CII (CP76) | Life cycle |
| ORF54 | 34758 | 35432 | 675 | + | Phage antirepressor protein KilAC domain | Life cycle |
| ORF55 | 35434 | 36609 | 1176 | + | Bacteriophage replication protein O | DNA metabolism and packaging |
| ORF56 | 36606 | 36767 | 162 | + | Hypothetical protein |  |
| ORF57 | 36788 | 37324 | 537 | + | Head completion protein Gp50 | Virion structure and assembly |
| ORF58 | 37325 | 37840 | 516 | + | Hypothetical protein |  |
| ORF59 | 37889 | 38356 | 468 | + | Putative cysteine dioxygenase | Auxiliary metabolic gene |
| ORF60 | 38353 | 38832 | 480 | + | Putative nuclease YbcO | DNA metabolism and packaging |
| ORF61 | 38829 | 39164 | 336 | + | gp4 | Virion structure and assembly |
| ORF62 | 39161 | 39409 | 249 | + | Hypothetical protein |  |
| ORF63 | 39820 | 40248 | 429 | + | Hypothetical protein |  |
